# Cancer In The News: Bias And Quality In Media Reporting Of Cancer Research

**DOI:** 10.1101/388488

**Authors:** Amanda Amberg, Darren N. Saunders

## Abstract

Cancer research in the news is often associated with sensationalising and inaccurate reporting, giving rise to false hopes and expectations. The role of study selection for cancer-related news stories is an important but less commonly acknowledged issue, as the outcomes of primary research are generally less reliable than those of meta-analyses and systematic reviews. Few studies have investigated the quality of research that makes the news and no previous analyses of the proportions of primary and secondary research in the news have been found in the literature. The main aim of this study was to investigate the nature and quality of cancer research covered in online news reports by four major news sources from USA, UK and Australia. We measured significant variation in reporting quality, and observed biases in many aspects of cancer research reporting, including the types of study selected for coverage, and in the spectrum of cancer types, gender of scientists, and geographical source of research represented. We discuss the implications of these finding for guiding accurate, contextual reporting of cancer research, which is critical in helping the public understand complex science and appreciate the outcomes of publicly funded research, avoid undermining trust in science, and assist informed decision-making.

## Introduction

Cancer is the second leading cause of death globally, accounting for 8.8 million deaths in 2015 [1]. The total annual economic cost of cancer was estimated at approximately US$ 1.16 trillion in 2010 [2] and it is estimated that 30-50% of cancer deaths could be prevented by modifying risk factors including tobacco exposure, alcohol consumption, obesity, exercise and infection [3]. Cancer is complex and challenging to study, and news reporting on cancer research is susceptible to hype, contradiction and misinformation. Clearly communicating the outcomes and context of research is key to helping non-specialists understand complex science, and assisting patients and families make informed decisions about modifying risk and treatment selection. Poor reporting practice may have serious consequences for public and scientific communities alike, including the generation of false or unmet expectations, potentially fuelling disappointment and a loss of trust in science [4, 5].

Few studies have focused on quantifying the types and quality of scientific research that gain attention in the news, which is arguably as important as accurate translation of research paper to news story. Even a news report that perfectly describes the findings of a study is of little value to the public if the findings themselves are based on weak evidence. Further, the reliability and long-term impact of primary research is rarely known at the time of publication. In cancer research and other fields, it has been demonstrated that initially reported effect sizes tend to be notably larger than those published in subsequent meta-analyses and that a striking proportion of publications will be refuted by follow-up studies [6-8]. This phenomenon was highlighted by Schoenfeld and Ioannidis in their meta-analysis on dietary risk and prevention of cancer [9]. News reporting on cancer research has previously been associated with poor accuracy, sensationalising headlines and presentation of conflicting information [10]. A preference towards reporting novel primary research stories with low replication likelihood often result in the refutation (or failed replication) of a research finding getting significantly less attention than the initial finding itself [11], reinforcing an ‘asymmetry of bullshit’ [12].

Previous analyses have observed that the distribution of news stories by cancer anatomical site mirrors incidence rates more closely than mortality rates but that certain cancer types were over- or under-represented. These distortions, potentially driven by personalisation bias (e.g., celebrity profiles) were also reflected in risk perception and discrepancies in funding for cancer research [13-15]. Others have also documented variation in quality, topic coverage and style of cancer research reporting in print media [16-18]. Previous research investigating the quality of translation from research papers and press releases to news stories highlights widespread problems with inadequate referencing and distorted reporting [19-24]. While the tabloid or ‘popular press’ was the main culprit, the same issues also exist in the more prestigious broadsheet news outlets [20, 25]. It may be tempting for researchers, journalists, philanthropic bodies and research institutions to sensationalise scientific findings in their pursuit of funding, readers or publicity. A common consequence of poor quality reporting is a hype cycle characterised by false expectations and subsequent disappointment [4]. Hype may be generated by journalists, institutional press releases, or the scientists themselves and can then be amplified through the media cycle [21, 26].

Primary research more easily lends itself to ‘breakthrough’ headlines since, by definition, it presents original data. Quality and reliability are not intrinsic features of meta-analyses and systematic reviews but depend on appropriate systematic methods [27, 28]. Nevertheless, the nature and purpose of these forms of secondary research – collating, comparing and re-analysing a set of primary studies to reduce uncertainty – render them less susceptible to error than individual primary studies [7]. Based on this assumption, secondary research is an important source of science news for the general public, yet there are indications of a reporting bias favouring primary studies [29]. In academic publishing, peer review and the accumulation of primary research papers followed by meta-analyses and review articles are designed to help filter inaccuracies of individual study outcomes but there is no equivalent formalised system in the news media beyond standard editorial oversight. Analysis of 734 front-page stories about medical research in major newspapers found that just over half were based on papers published in peer-reviewed journals, a minority of these being systematic reviews [29]. This emphasises the need for better characterisation of the types and quality of cancer research studies that gain attention in the news.

Hence, this study aimed to analyse selection bias and quality of cancer research published in four major news sources from the UK, USA and Australia over a six-month period in 2017.

## Methods

### News report collection

Twenty news reports were sampled from each of the online versions (published between March and September 2017) of *The Guardian* (UK edition), The *New York Times* (*NY Times*), *The Sydney Morning Herald* (*SMH*) and the *Australian Broadcasting Corporation* (*ABC*), generating a total dataset of 80 reports. The following search terms were used within the search function on each source’s website: *‘cancer study’, ‘cancer research’, ‘cancer science’, ‘targeted therapy’, ‘cancer screening’, ‘cancer screening study’, ‘tumour research’, ‘tumor research’, ‘cancer treatment’, ‘cancer genetics’* and *‘cancer scientists’*. Only original reports were included in the sample, reports re-published from other news sources were excluded to avoid overlap in the data. The reports had to discuss a study investigating a cancer-related issue including epidemiology, carcinogens, screening, diagnostics, therapies, basic biology, risk or prevention. General reports which dealt with cancer-related topics but which did not base the discussion on a specific study were excluded. When several studies were discussed and no one central study could be distinguished, the report was excluded. Details of individual reports (including Pubmed ID (PMID), quality scores and hyperlinks to reports) are contained in Supplementary Table 1.

### Classification and scoring

Where possible, the study discussed in each media report was traced and classified as basic research, clinical research, epidemiological research, meta-analysis or systematic review according to the classification of Röhrig *et al*. [30]. Original research sources cited in news reports were classified as either published paper, conference, report, press release, pre-print article, funding source/researcher or unknown. News report quality was scored according to a matrix based on eleven criteria adapted from previous studies (Table 1) [19, 21]. Australian cancer incidence and mortality statistics were obtained from the Australian Institute of Health and Welfare [31]. Gender of senior authors (i.e. names appearing in first and last position on author list) and quoted experts in each report was also quantified using pronouns quoted in reports or on homepage. Source nationality was classified according to primary academic affiliation of corresponding authors.

**Table 1.**
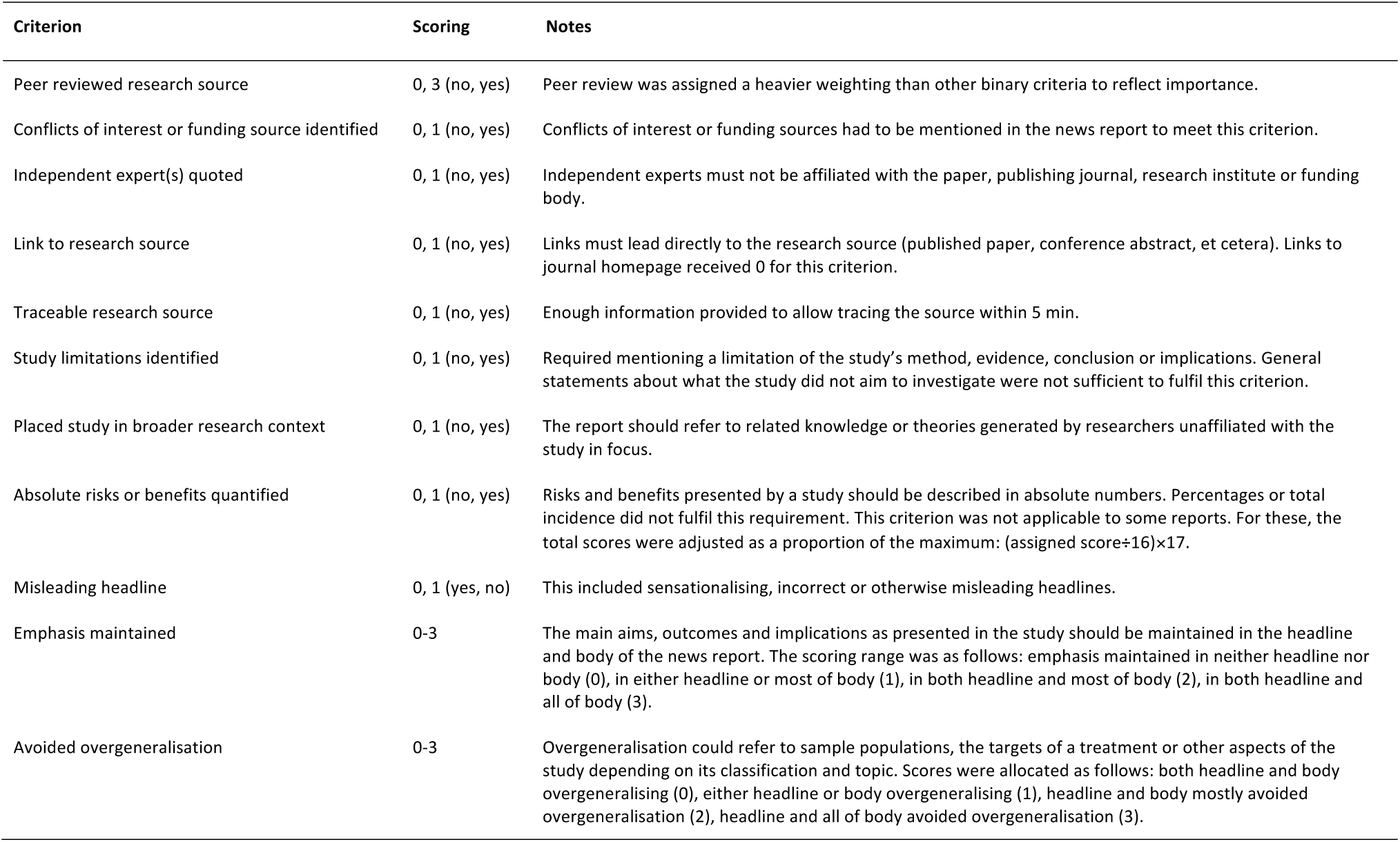
News Report Quality Scoring Criteria.

### Data analysis

Chi-square or Fischer’s exact tests were used to compare the categorical variables (primary or secondary study), assuming statistical significance if P<0.05. Quality scores were aggregated according to Table 1 and mean scores calculated. Statistical significance of differences in mean scores between the news sources was tested using one-way ANOVA, assuming Gaussian distribution, for multiple comparisons with a Tukey test. When analysing national bias, all studies that had been conducted as international collaborations were included in the ‘international’ category.

## Results

### Content bias

The long-term implications of primary research findings are rarely known at the time of publication. Indeed, while basic research is absolutely essential to scientific progress, there is evidence that a remarkable proportion of published results will be refuted by further investigation or that subsequent meta-analyses will report notably smaller effect sizes, in cancer as well as other research areas [6-8]. Schoenfeld and Ioannidis [9] published a meta-analysis highlighting this phenomenon in research on cancer risk and prevention. Similar trends have been observed in basic medical research, where only a small fraction of the most encouraging early findings end up in clinical use [32]. This becomes important when scientific research papers gain attention in the mainstream press, the predominant source of science news for the public. However, there is limited literature on content bias in science news reporting. Quality and style have been shown to vary across news outlets [16, 17], but even the largest newspapers with the best overall standards tend to cover more studies with poorer methodology and observational studies over RCTs or systematic reviews [33-35].

To investigate the distribution of different study types in cancer research reporting, we quantified the number of primary and secondary studies and their subtypes in a sample of 80 news stories from 4 different outlets. Of the samples reports, 92.5% (74/80) were based on primary research studies (Fig. 1a). When studies were further classified by subtype, epidemiological studies were the most prevalent overall, accounting for 38.75% of reports (31/80), followed by clinical and basic research at 28.75% (23/80) and 23.75% (19/80) respectively (Fig. 1b). Secondary studies consisted of four systematic reviews and two meta-analyses (Fig. 1b). One study did not fit in any of the categories and was therefore classified as ‘other’.

**Figure 1.**
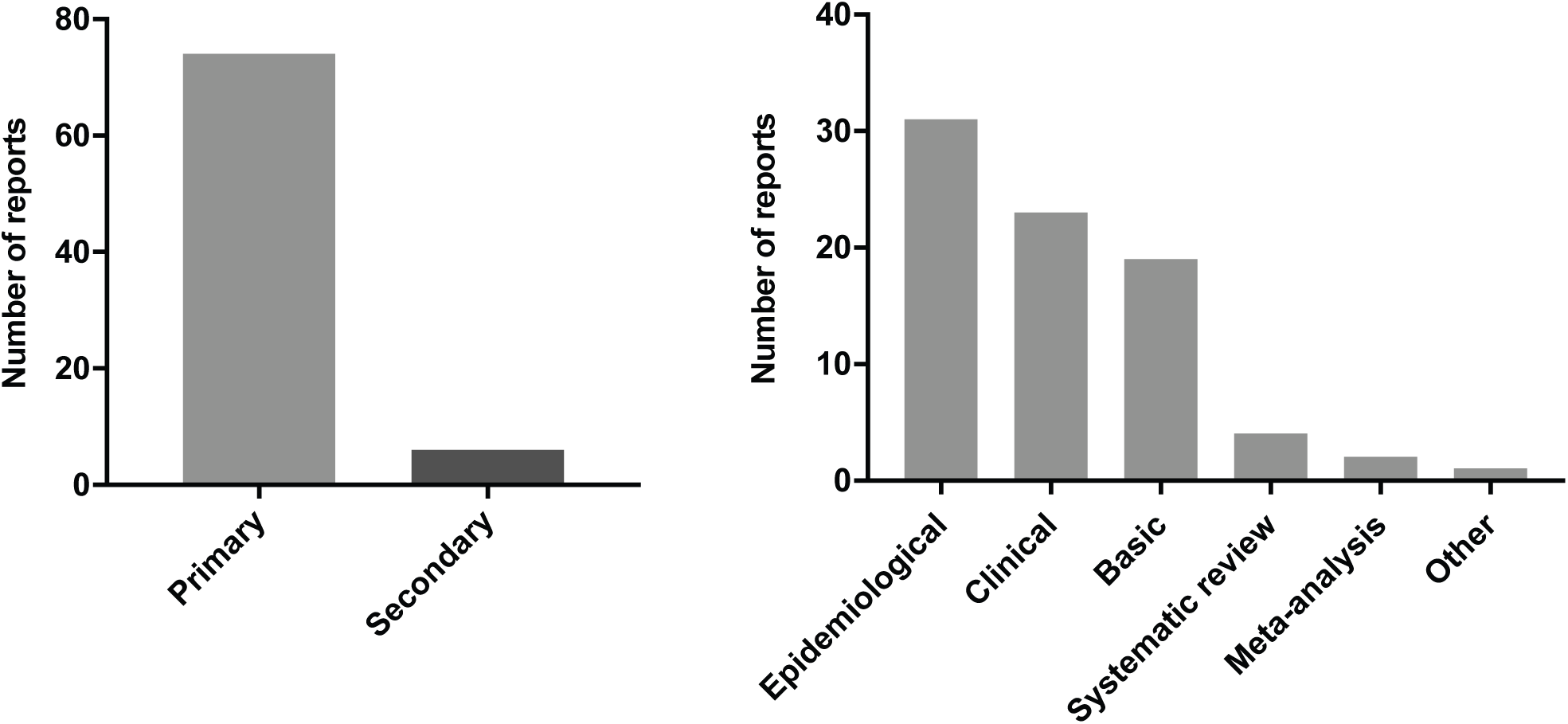
Study type bias represented in online news reports about cancer: **A.** Frequency of primary and secondary research represented in online news reports about cancer research. **B.** Distribution of research subtypes represented in news reports (n=80).

We next classified and quantified the source of research studies cited in our sample of 80 news reports. Most news reports (68.75% or 55/80) were based on peer-reviewed papers (Fig. 2), sometimes accompanied by a press release. Research published outside of traditional academic journals (e.g. reports issued by government agencies) were the basis of 12.5% (10/80) of news items, and 10% (8/80) of reports were based on conference presentations (five of these were published concurrently with a major annual cancer conference). Two news reports were based on institutional press releases without an associated published scientific paper, and two others referred only to the researcher or funding source. The source of three news reports could not be traced (Fig. 2).

**Figure 2.**
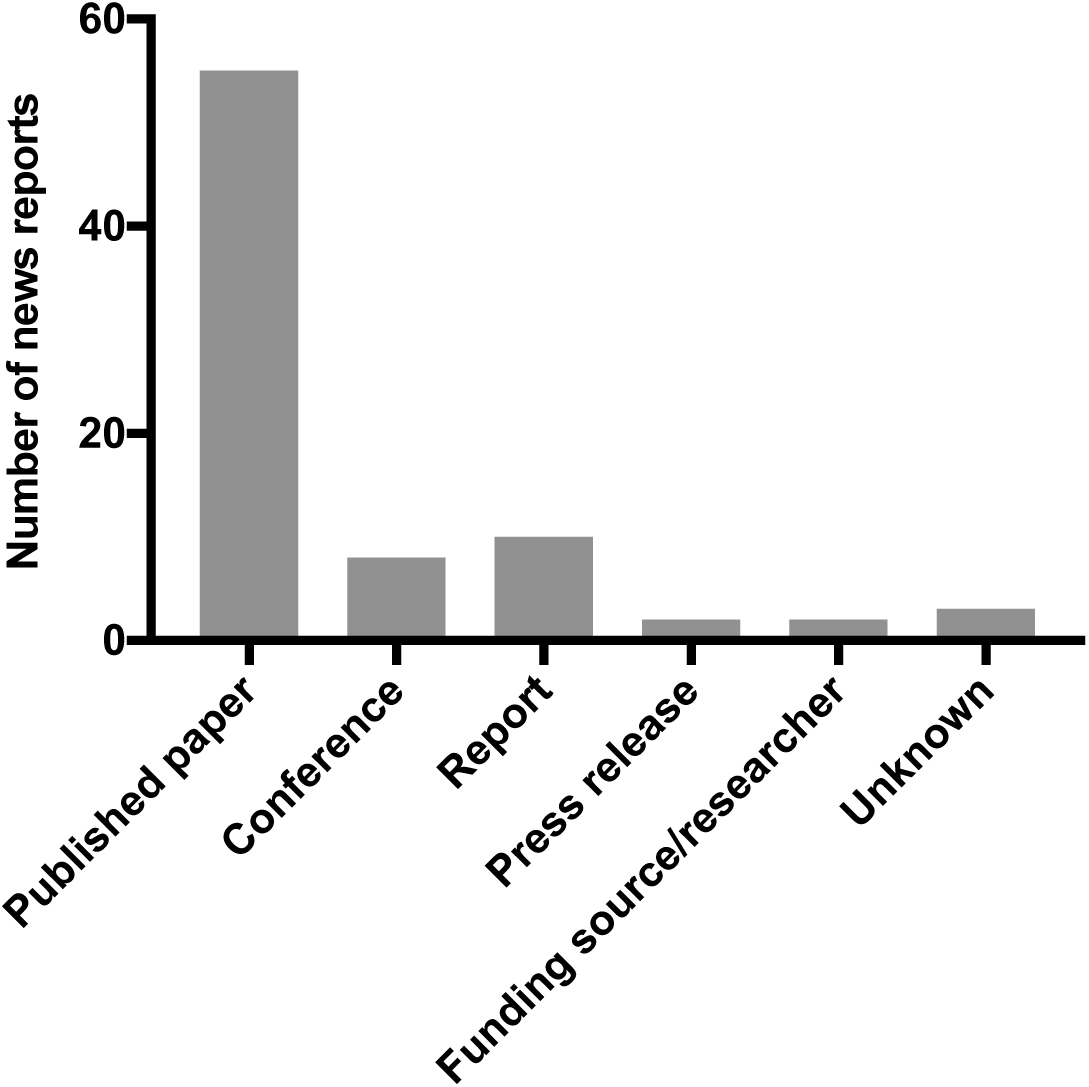
Distribution of research types forming the basis of online news reports about cancer research (n=80).

### Cancer types represented in reporting

We examined the distribution of cancer types (defined by anatomical site) represented in our cohort of 80 news reports. The most frequent category observed was non-specific (i.e. not related to a specific cancer type), representing 18/80 (22.5%) reports and possibly reflecting a strong bias towards more basic research on disease mechanisms and risk. Among cancer types explicitly identified in news reports, breast (15%), melanoma (11.3%), lung (8.8%) and blood (8.8%) cancers were the most frequently reported (Figure 3a). Reports specifically mentioning gastric, testicular, brain and pancreas cancer were the least frequently observed, with each only being represented in a single report. Many cancer types were not represented in news reports at all during sample period. When analysed in the context of relative rates of incidence and mortality of specific cancer types, we observed a strong correlation between reporting and incidence of specific cancer types (R^2^ = 0.594, p=0.0013) but not with mortality (Figure 3b,c). Research on cervical cancer was reported more frequently than would be expected relative to incidence, while prostate and colorectal cancer were under-represented in news reports (Figure 3b). Relative to mortality, cervical, melanoma and breast cancer were over-represented, while lung, pancreas, and colorectal cancer were under-represented (Figure 3c).

**Fig 3:**
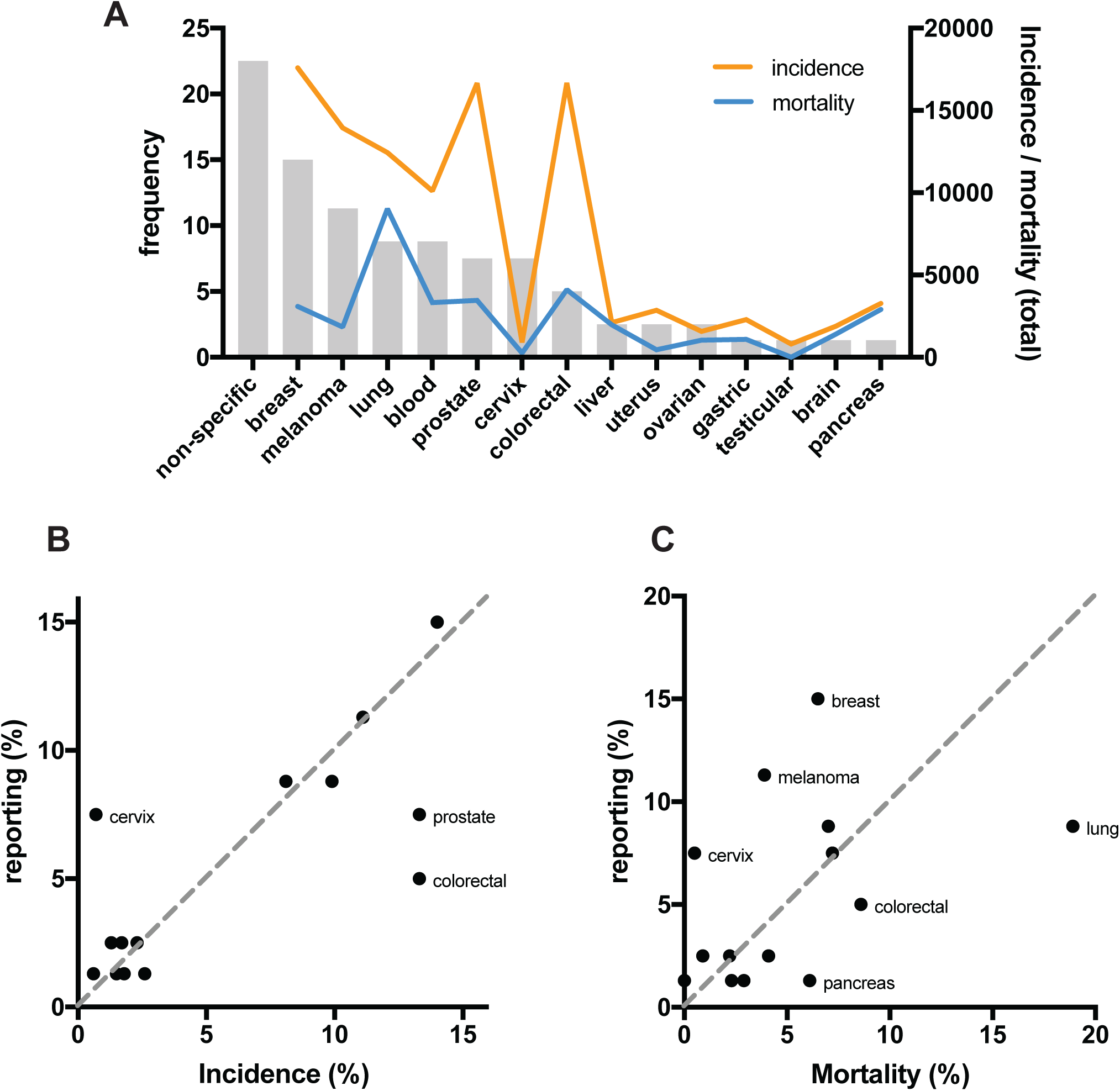
Analysis of bias in cancer types reported. **A.** Distribution of cancer types in research studies covered in news reports, with Australian incidence and mortality rates; Reporting frequency (as a proportion of total) compared with relative incidence **(B)** and mortality **(C).**

### Reporting quality

News reporting of research is of most value to readers if it accurately conveys the outcomes, context and implications of the research. Particularly in the context of serious diseases like cancer, accurate reporting is critical for informing decisions on modifying risk, choice of intervention, understanding prognosis, etc. The level of consensus between news articles and related original research papers as a marker of accuracy has been the focus of extensive research. Almost three decades ago, Singer [19] proposed an analytical model for scoring accuracy with eleven criteria, including issues of incorrect statements, misleading headlines, sensationalisation and overgeneralisation, and these have become benchmarks for evaluating quality of media reports on research. Modified versions of Singer’s method have revealed poor citation practice and frequent misleading reporting in the news [22-24]. Although the tabloid press is the biggest culprit in misinterpreting science news, studies have demonstrated that a lack of quality also exists in broadsheet newspapers [20, 25]. Today’s online environment is thought to result in many major news outlets utilising the same sources of information, potentially resulting in an amplified spread of poor quality reporting [36]. Research institutes and funding bodies seeking publicity and philanthropic support may exploit this space as press releases play a major role in shaping the content of many news articles [21].

We measured reporting quality in our cohort using a scoring matrix modified from Singer [19] and Taylor [21] for comparison between various news outlets (Table 1). The *NY Times* had the highest average quality score (12.9), while the lowest average scores were seen in the Australian news sources (9.5 and 8.8 for *ABC* and *SMH* respectively), with reports in *The Guardian* averaging a quality score of 11.2 (Fig. 4). Reports in the *NY Times* also displayed the most consistent quality scores, although there was variability in quality observed in all news sources. Where online readership statistics were available (i.e. for the 20 studies published by ABC), we observed no relationship between reporting quality and readership (not shown). Similarly, we observed no obvious relationship between study type and reporting quality, but the low number of secondary studies reported limited this analysis.

**Figure 4.**
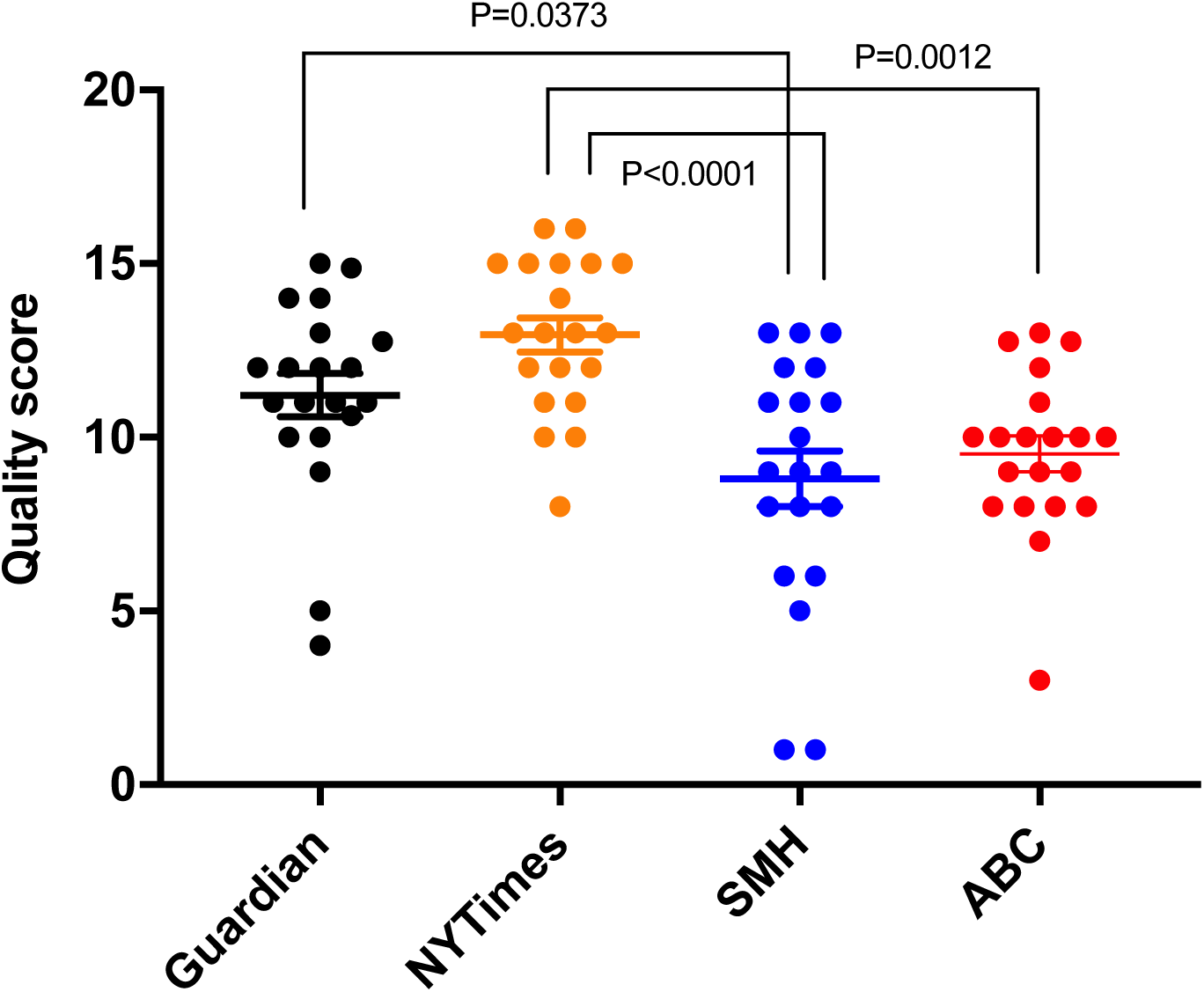
Quality scores of news reports on cancer research. Each point represents the score of an individual news report. Bars indicate the mean ± SEM for each news outlet (n=20 for each).

### National bias in reporting of cancer research

While research performed in the USA dominates scientific output in terms of papers published [37], reporting on research in a local context has important implications for both consumers and scientists alike. Different risk factors may have proportionally different significance for various audiences and it is critical that scientists are able to reach the most relevant audiences on topics of local importance. For example, understanding the role of UV exposure in melanoma risk is an important consideration in Australia. Further, it is important for both scientists and consumers alike to have outcomes of publicly funded research communicated to taxpayers.

To analyse national bias in cancer research reporting, we analysed the country of origin of research cited in each news report - determined by the primary affiliation of the corresponding author of the research publication (where available). At least half of the reports in each news source were based on research from the same country in which the news organisation was based (Fig. 5). *The Guardian* (UK edition) had the most diverse national origin of research cited, with only 50% of reports based on research performed primarily in the UK. In contrast, 70% of reports in the *NY Times* were based on research studies performed primarily in the US and 72.5% of Australian news reports were based on Australian research (65% and 80% by *SMH* and *ABC*, respectively) (Fig. 5). Viewed from the opposite perspective, the great majority (29/30 studies, or 97%) of Australian research studies represented in our cohort were only reported in Australian news outlets. In other words, only 1/30 Australian studies received international coverage (Fig 5).

**Figure 5.**
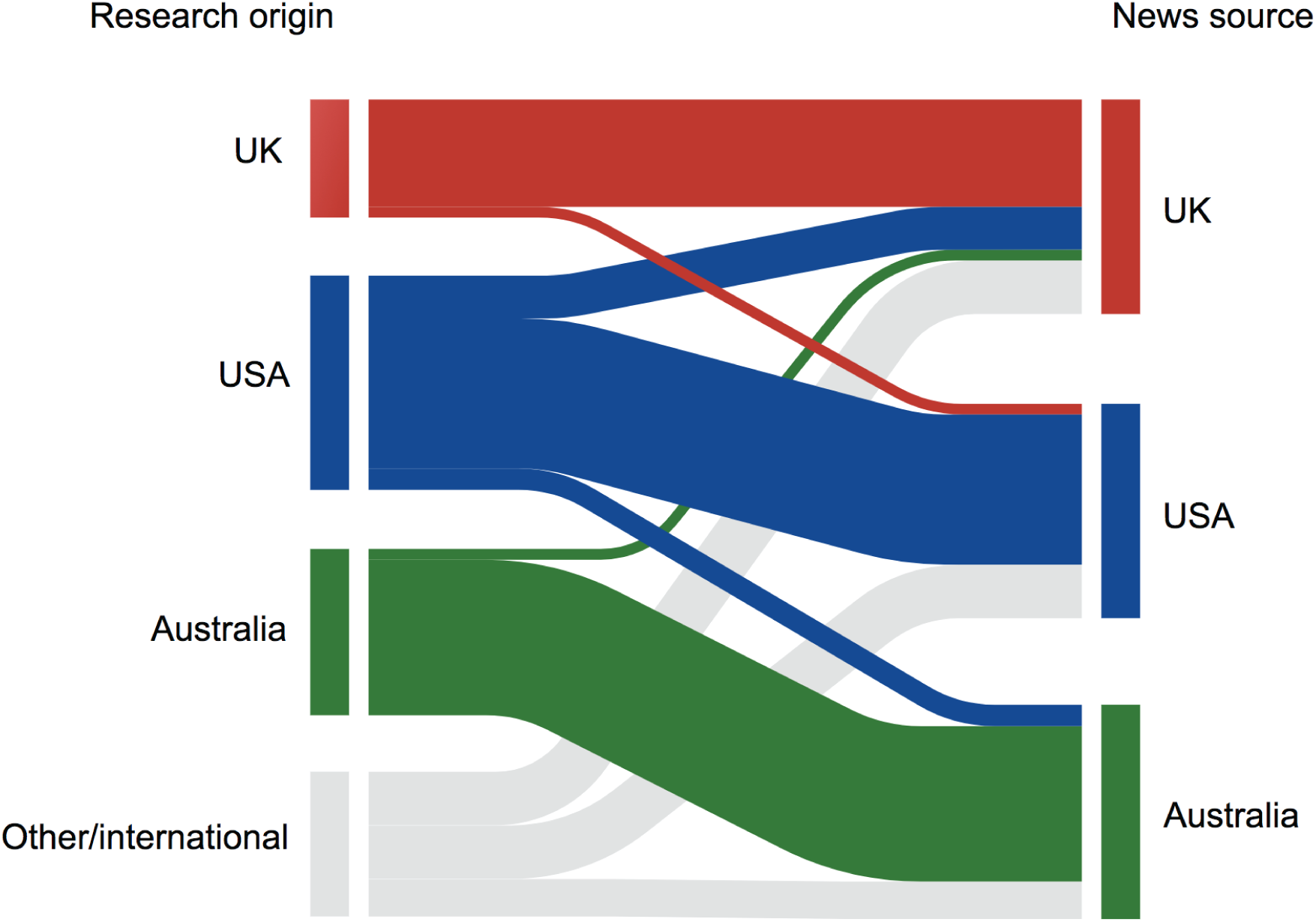
National bias in reporting of cancer research. Sankey chart showing relationships between country of research origin (left) to country in which reporting news organisation is based (right), with bar sizes representing proportional representation. The UK is represented by *The Guardian*, USA by *NY Times* and Australia by *SMH* and *ABC*.

### Gender bias in reporting of cancer research

Female scientists face a suite of documented biases [38-41], and a number of studies have established that women are under-represented in news media [42, 43]. More specifically, the systematic under-recognition of female scientists – known as the *Matilda effect* [44] – has been demonstrated in science communication, where publications from male authors are associated with greater perceived scientific quality [45]. Across our entire cohort of 80 news reports, we observed a significant gender bias among senior authors, with 60% (67/112) of research studies reported having male senior authors (Fig 6a). We also observed a significant bias toward male experts being quoted in news reports, with 68% (100/148) of quoted experts being male. A similar trend was observed in individual news outlets, with the exception of the *ABC* - where equal representation of male and female senior authors was observed in the studies forming the basis of news reports (Fig 6b). The bias towards quoting male experts in online reports about cancer research was consistent across individual news outlets (Fig 6c).

**Figure 6.**
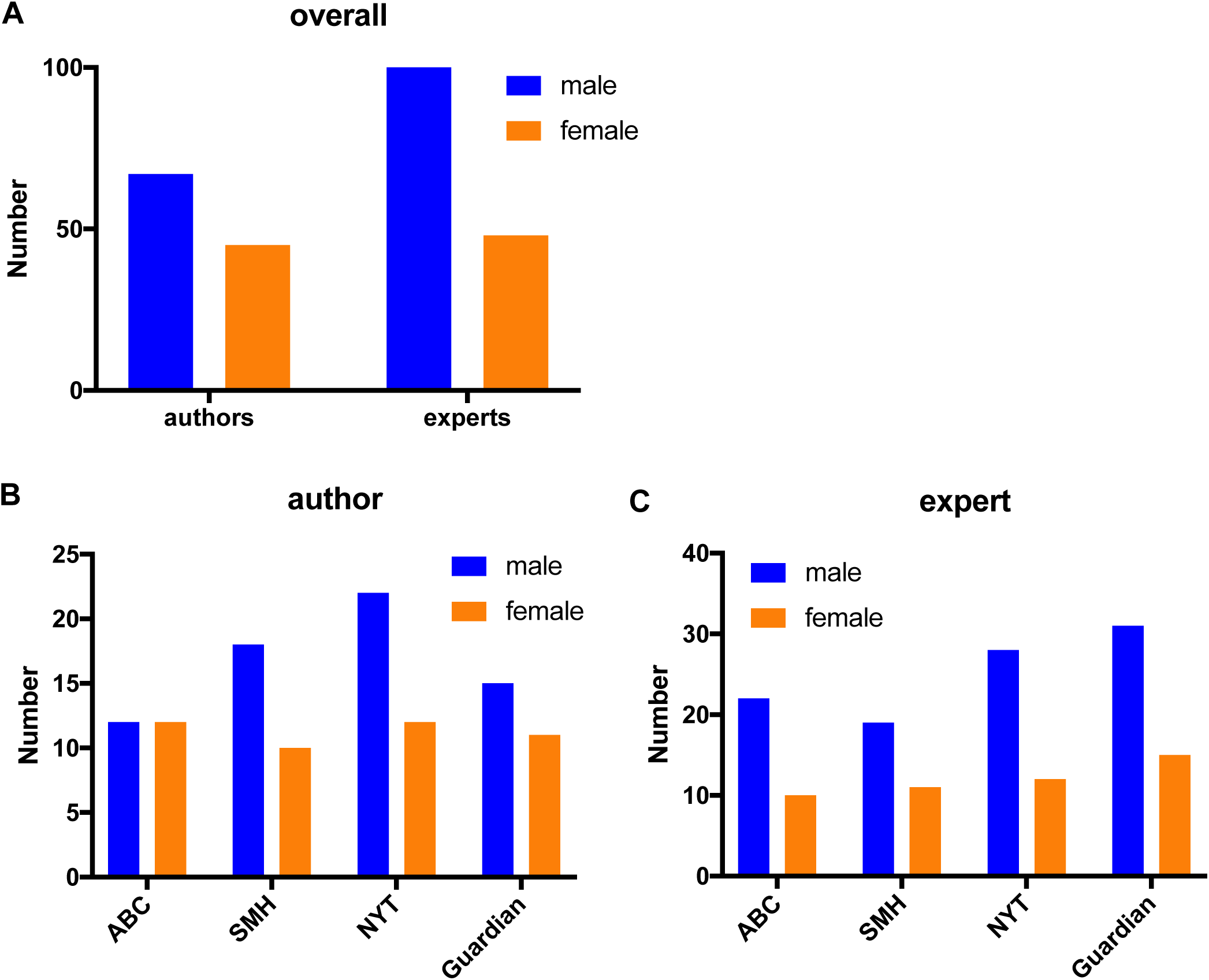
Gender bias in senior authors and quoted experts in news reports. Distribution of senior authors and experts across the entire cohort (n=80), and distribution of authors **(B)** and experts **(C)** in individual news outlets (n=20 each).

## Discussion

Clear communication of research outcomes in context is important in helping non-specialists understand the often complex and challenging contents of scientific publications. Poor reporting may hinder informed decision-making about modifiable risks and treatment selection, generate false or unmet expectations, and undermine trust in science [4, 5]. In order to better understand factors influencing reporting of cancer research we analysed the distribution of study types, research sources, reporting quality, gender bias, and national bias in online news reports by four major news outlets in USA, UK and Australia over a six-month period in 2017.

Our analysis demonstrated that primary research studies predominate over secondary studies. This may reflect a similar trend in the publication frequency of these study types – by their nature, systematic reviews and meta-analyses are less abundant than primary studies – or the way institutes promote their research. Further, evidence of a skew towards lower quality studies in the news [33] indicates that publication bias might be an underlying factor in newspapers favouring research with poorer methodology. Similar trends have been observed in other publications, with observational studies on average receiving more attention than both systematic reviews and randomised controlled trials (RCTs) [34, 35]. This is problematic as the features thus attracting media attention are often a key driver of hype, confusion and false expectations. Hence, our observed bias towards reporting primary studies may increase the risk of generating hype about cancer research via news reporting.

When searching for novel stories, journalists are likely to favour primary research findings due to their novelty and frequently larger effect sizes [7] but basic research articles are also the most susceptible to sensationalisation [36]. While experienced scientists and many journalists likely know to view these papers as potentially useful pieces in a much greater puzzle, the general population may not have the experience or specialist knowledge to interpret individual reports critically in a broader context. Despite blame commonly being attributed to journalists and press releases, statements in the research articles themselves are often exaggerated and may generate hype. For example, many observational studies in high impact journals contain advice on clinical practice without mentioning the need for confirmation by RCTs [26] or fail to discuss population biases, small samples sizes, or difficulties in translating animal studies to humans.

The low occurrence of secondary studies reported in our sample of online reports is consistent with previous findings showing that systematic reviews represented a small proportion of medical research news [29]. This trend is likely reflected in the frequency of these studies published in journals, although we could not find any literature that directly investigated the publication frequency of different study types in medical research. We found a majority of online news articles reporting on peer-reviewed papers, however this may be partly explained by our exclusion of more general news articles that did not report on a specified study. Previous studies have highlighted inconsistent quality and accuracy of science news reporting practices at multiple levels, ranging from institutional press releases to news pieces [16, 17, 21], and found that study types with poorer methodology gain more media coverage than research based on stronger evidence [33-35]. Our analysis of reporting quality and study type distribution in online news is consistent with previous evidence of poor quality reporting by broadsheet news sources [19, 20, 23] and a bias towards primary research [6, 9, 46, 47]. This predominance of primary studies in the news increases the likelihood that the general public is left with a distorted perception of cancer research and an inaccurate view of scientific progress [36]. This is partly due to the increased risk of premature statements about yet un-replicated research outcomes and the greater likelihood of consequent contradiction or refutation. For example, cancer drug attrition rates are characteristically high, with only 5% of preclinical anticancer drugs reaching clinical use, and reproducibility of high-impact studies in this area is low [46]. Even therapies tested in phase II clinical trials rarely succeed in reaching clinical practice [47].

Reporting quality scores varied within and across the news sources. Mean quality scores indicated similarity between *The Guardian* and *NY Times* but were significantly lower in the Australian news sources. Pointing to areas for potential future improvement, *SMH* and *ABC* often failed to consult an independent expert, provide a link to the study being discussed, maintain emphasis, and avoid overgeneralisation. While all sources regularly failed to mention limitations of the study being reported, Australian news more often omitted such statements. It should also be noted that availability of resources in different media outlets may have a significant influence on reporting quality but we were not able to measure this effect. The online news media’s selection of research sources has previously been evaluated by testing whether *Altmetric* scores correlate to citation indices or journal impact factors, with conflicting results [48, 49]. Authors have argued that this shows that the best research articles are not necessarily the ones that make it to the news. While this may be true, methods based on *Altmetric* scores will not capture the quantity of news articles that discuss research without directly referencing a study. Furthermore, other studies have indicated that alternative metrics may generate inconsistent results depending on data collection methods [50], and that citation index is not necessarily the best marker of study quality [51, 52]. We did not observe any relationship between quality and readership in reports by the *ABC* but unfortunately could not expand this analysis as audience data was not available for other news outlets. The low number of secondary studies made the cohort in this study underpowered for an analysis comparing type and quality.

Previous analysis of cancer research stories on the *BBC* website from 1998-2006 found a heavy focus on breast cancer, followed by lung and prostate cancers [53]. Almost a quarter of news reports in our cohort did not refer to a specific cancer type, possibly reflecting the strong bias towards reporting of basic research on disease mechanisms and risk. However, an important underlying factor in this trend may also be the emerging trend to define cancer type by molecular rather than anatomical classifiers [54, 55]. We observed a correlation between the representation of various cancer types (classified by anatomical site) and the relative incidence rates of those cancer types (Figure 3b), but no relationship between reporting frequency and relative mortality rates (Figure 3c). Cervical, melanoma and breast cancer were over-represented relative to their respective mortality rates, while lung, pancreas, and colorectal cancer were under-represented. Continued public and media interest on the effectiveness of the HPV vaccine *Gardasil*, and changes in screening practices may be a relevant consideration in the over-representation of reports focussing on cervical cancer. Over or under representation of different cancer types in research reporting can skew public awareness of risk factors and may also drive inequities in public and philanthropic funding of research directed as specific cancer types.

We observed a striking national bias, where news outlets were more likely to report on research performed in the same country they were based. If the distribution of studies reported in all news outlets mirrored global research output, the vast majority of reports would be based on studies performed in the US, as they dominate in terms of papers published [37]. However, a perfect reflection of global output is not necessarily desirable. It may be difficult for local scientists to get media attention outside their own country, and institutional press offices may have stronger contact networks with local reporters. Reporting research of relevance to distinct geographical areas (e.g. epidemiological investigations on specific populations) may be very important in informing the public with regards to local risk factors and outcomes of publicly funded research. Hence, the predominance of epidemiological studies in news reports is one possible contributor to the observed national bias. Conversely, prioritising reporting on local research means the public may not get access to important information from broader sources, although the modern online environment puts global news within reach of the majority of people.

The *Matilda effect*, which describes the systematic under-recognition of female scientists, has been extended to science communication, where greater perceived scientific quality is associated with publications from male authors [44, 45]. Further, under-representation of women in news media more generally is well established [42, 43]. We observed a striking gender bias in both study selection and reporting in our cohort, with the majority of reports based on studies with male senior authors and quoting male experts. This bias has potential to compromise high-quality coverage of research by limiting diversity of opinion, and likely serves to reinforcing stereotypes and further entrench gender inequity among researchers by providing public visibility and recognition predominantly to male scientists. The suite of biases faced by female scientists is well documented [38-41], and the gender bias in news reports likely reflects an underlying predominance of men among the ranks of senior scientists. The possibility of other underlying biases can’t be excluded, including differences in the availability and/or willingness of male experts to speak to journalists. Regardless, our data highlights a need for journalists, scientists and institutes to significantly improve efforts to ensure equal representation of male and female scientists in news reports on cancer research.

### Limitations

As cancer research output is not expected to fluctuate notably throughout the year, it was assumed that a six-month sample period limited to 20 reports from each of the four news outlets would provide a representative cohort to analyse. An exception was conference-based news reports, which peaked at the time of a major cancer conference in June (ASCO annual meeting, https://am.asco.org/). In all news outlets apart from *The Guardian*, the 20 reports comprised the majority of relevant reports within the selected time frame and should thus be considered representative samples of each source. Although the size of our cohort limits a comprehensive representation of national reporting trends, the chosen sources are all major news outlets in their respective countries and so likely provide a reasonable indication of broader trends. It was also assumed that all studies could be classified according to the model outlined by Röhrig et al. [56]. Secondary studies were too rare to allow analysis of any potential relationship between study type and reporting quality, which should be investigated in larger datasets.

### Recommendations and conclusions

The quality indicators measured in our cohort provide useful guidelines for journalists to consider in providing the most informative and accurate reporting of research. These are particularly relevant to minimising the potential for hyperbole, providing an objective account of research outcomes and implications, and in assisting readers to critically assess the report. For example, reports in the Australian news outlets (*SMH* and *ABC)* often failed to consult an independent expert, provide a link to the research study, and avoid overgeneralisation. While all sources regularly failed to mention limitations of the study being discussed, Australian news more often omitted such statements. These limitations may reflect limited time and/or resources of reporters or a trend towards having fewer specialist reporters in Australian media, but we are unable to quantify these in our dataset. Our data highlight the importance of considering which research types are selected for coverage. Acknowledging the importance of journalists providing independent and novel context, interpretation and insight to individual stories, a better representation of the current state in cancer research would be achieved by attempting to cover a more balanced proportion of primary and secondary studies, from national and international sources. Scientists and journalists should also take care to mention the limitations of novel ideas in research and refrain from presenting findings as certain.

Conversely, when communicating with the news media, scientists should be conscious of the possible discrepancy between impact in the scientific community and among the general public. Limitations and uncertainties should always be highlighted. Where possible, readers should be made aware of what type of study is being reported on, whether it is peer-reviewed and how strong the supporting evidence is. These are important considerations, as even prestigious newspapers do not necessarily give a fair representation of current progress in cancer research. As far as we know, this is the first combined analysis of study type distribution, reporting quality, and other biases in cancer research reporting. These data highlight the presence of significant biases and provide a basis for improving the selection of studies being selected for media coverage, and the way those studies are reported. Future analyses should build on the findings reported here by incorporating the long-term outcomes and impact of the studies that appear in the news media. It would also be useful to evaluate to what extent corrections follow in the news after one of these studies have been refuted or a declared ‘breakthrough drug’ fails to reach the market. Further, analyses of relationships between readership, study type and reporting quality would offer insight into how demand-driven these biases may be.

Accurate, contextual reporting of cancer research is imperative in helping the public understand complex and challenging science and appreciate the outcomes of publicly funded research, avoid undermining trust in science, and assist informed decision-making.

## Acknowledgments

The authors wish to thank Dr Adam Dunn and Dr Jonathon Webb for thoughtful and constructive discussion and comments on draft mansucripts.

